# A systematic survey of centrality measures for protein-protein interaction networks

**DOI:** 10.1101/149492

**Authors:** Minoo Ashtiani, Ali Salehzadeh-Yazdi, Zahra Razaghi-Moghadam, Holger Hennig, Olaf Wolkenhauer, Mehdi Mirzaie, Mohieddin Jafari

**Author notes:** These authors have contributed equally to this work (Minoo Ashtiani, Ali Salehzadeh-Yazdi). Corresponding Authors: Mehdi Mirzaie, Mohieddin Jafari www.jafarilab-pasteur.com.

## Abstract

**Background:** Numerous centrality measures have been introduced to identify “central” nodes in large networks. The availability of a wide range of measures for ranking influential nodes leaves the user to decide which measure may best suit the analysis of a given network. The choice of a suitable measure is furthermore complicated by the impact of the network topology on ranking influential nodes by centrality measures. To approach this problem systematically, we examined the centrality profile of nodes of yeast protein-protein interaction networks (PPINs) in order to detect which centrality measure is succeeding in predicting influential proteins. We studied how different topological network features are reflected in a large set of commonly used centrality measures.

**Results:** We used yeast PPINs to compare 27 common of centrality measures. The measures characterize and assort influential nodes of the networks. We applied principal component analysis (PCA) and hierarchical clustering and found that the most informative measures depend on the network’s topology. Interestingly, some measures had a high level of contribution in comparison to others in all PPINs, namely Latora closeness, Decay, Lin, Freeman closeness, Diffusion, Residual closeness and Average distance centralities.

**Conclusions:** The choice of a suitable set of centrality measures is crucial for inferring important functional properties of a network. We concluded that undertaking data reduction using unsupervised machine learning methods helps to choose appropriate variables (centrality measures). Hence, we proposed identifying the contribution proportions of the centrality measures with PCA as a prerequisite step of network analysis before inferring functional consequences, e.g., essentiality of a node.

## Introduction

Essential proteins play critical roles in cell processes such as development and survival. Deletion of essential proteins is more likely to be lethal than deletion of non-essential proteins (1). Identifying essential proteins conventionally had been carried out with experimental methods which are time-consuming and expensive, and such experimental approaches are not always feasible. Analyzing high-throughput data with computational methods promises to overcome these limitations. Various computational methods have been proposed to predict and prioritize influential nodes (e.g. proteins) among biological networks. Network-based ranking (i.e. centrality analysis) of biological components has been widely used to find influential nodes in large networks, with applications in biomarker discovery, drug design and drug repurposing (2). Not only in molecular biology networks but also in all types of networks, finding the influential nodes is the chief question of centrality analysis (3). Examples include predicting the details of information controlling or disease spreading within a specific network in order to delineate how to effectively implement target marketing or preventive healthcare (4-6). Several centralities measures (mostly in the context of social network analyses) have been described (3) in the last decades. A comprehensive list of centrality measures and software resources can be found on the CentiServer (7).

The correlation of lethality and essentiality with different centrality measures has been subject of active research in biological areas, which has led to the centrality-lethality rule (1). Typically, some classic centrality measures such as Degree, Closeness, and Betweenness centralities have been utilized to identify influential nodes in biological networks (5). For example, in a pioneering work, the authors found that proteins with the high Degree centrality (hubs) among a yeast PPIN is likely to be associated with essential proteins (1). In another study, this rule was re-examined in three distinct PPINs of three species which confirmed the essentiality of highly connected proteins for survival (8). Similar results were reported for gene co-expression networks of three different species (9) and for metabolic network of *Escherichia coli* (10, 11). Ernesto Estrada generalized this rule to six other centrality measures. He showed that the Subgraph centrality measure scored best compared to classic measures to find influential proteins, and generally using these measures performed significantly better than a random selection (12). However, He and Zhang showed that the relationship between hub nodes and essentiality is not related to the network architecture (13). Furthermore, regarding the modular structure of PPINs, Joy *et al.* concluded that the Betweenness centrality is more likely to be essential than the Degree centrality (14). The predictive power of Betweenness as a topological characteristic was also mentioned in mammalian transcriptional regulatory networks which was clearly correlated to Degree (15). Recently, it has been shown that presence of hubs, i.e. high Degree centralities, do not have a direct relationship with prognostic genes across cancer types (16).

On the other hand, Tew and Li demonstrated functional centrality and showed that functional centrality correlates more strongly than pure topological centrality (17). More recently, Peng *et al.* introduced localization-specific centrality and claimed that their measure is more likely to be essential in different species (18). Khuri and Wuchty introduced minimum dominating sets of PPIN which are enriched by essential proteins. They described that there is a positive correlation between Degree of proteins in these sets and lethality (19). In these studies, the solution of the controversy is ascribed to utilizing biological information.

Similar in methodology but different in the underlying physical system that the network represents, some other studies attempted to quantify correlations between several classic centrality measures. In 2004, Koschützki and Schreiber compared five centrality measures in two biological networks and showed different patterns of correlations between centralities. They generally concluded that all Degree, Eccentrecity, Closeness, random walk Betweenness and Bonacich’s Eigenvector centralities should be considered to find central nodes and could be useful in various applications without explaining any preference among them (20). Two years later, they re-expressed pervious outcomes by explaining the independence behavior of centrality measures in a PPIN using 3D parallel coordinates, orbit-based and hierarchy-based comparison (21). Valente *et al.* examined the correlation between the symmetric and directed versions of four measures which are commonly used by the network analysts. By comparing 58 different social networks, they concluded that network data collection methods change the correlation between the measures and these measures show distinct trends (22). Batool and Niazi also studied three social, ecological and biological neural networks and they concluded the correlation between Closeness-Eccentricity and Degree-Eigenvector and insignificant pattern of Betweenness. They also demonstrated that Eccentricity and Eigenvector measures are better to identify influential nodes (23). In 2015, Cong Li *et al.* further investigated the question of correlation between centrality measures and introduced a modified centrality measure called *m*th-order degree mass. They observed a strong linear correlation between the Degree, Betweenness and Leverage centrality measures within both real and random networks (24).

However, there is no benchmark for network biologists that provides insight, which of the centrality measures is suited best for the analysis of the given network. The result of the centrality analysis of a network may depend on the centrality measure used which can lead to inconsistent outcomes. Previously, a detailed study showed that the predictive power and shortcomings of centrality measures are not satisfactory in various studies (25). While these centrality measures have proven to be essential in understanding of the roles of nodes which led to outstanding contributions to the analysis of biological networks, choosing the appropriate measure for given networks is still an open question. Which measure identifies best the centers of real networks? Do all measures independently highlight the central network elements and encompass independent information or are the measures correlated? Is the computation of all these measures meaningful in all different networks or does the best measure depend on the network topology and the logic of the network reconstruction? In this study, we used unsupervised machine learning to compare how well the most common centrality measures characterize nodes in networks. We comprehensively compared 27 distinct centrality measures applied to 14 small to large biological and random networks. All biological networks were PPINs of the same set of proteins which are reconstructed using a variety of computational and experimental methods. We demonstrated how the ranking of nodes depends on the network structure (topology) and why this network concept i.e. centrality deserves renewed attention.

### Materials and methods

The workflow of this study was schematically presented in Fig. 1. Our workflow started by constructing and retrieving networks, followed by global network analysis. The centrality analysis and comparing them using machine learning methods were the next main steps. See basic definitions for more details.

**Fig. 1.**
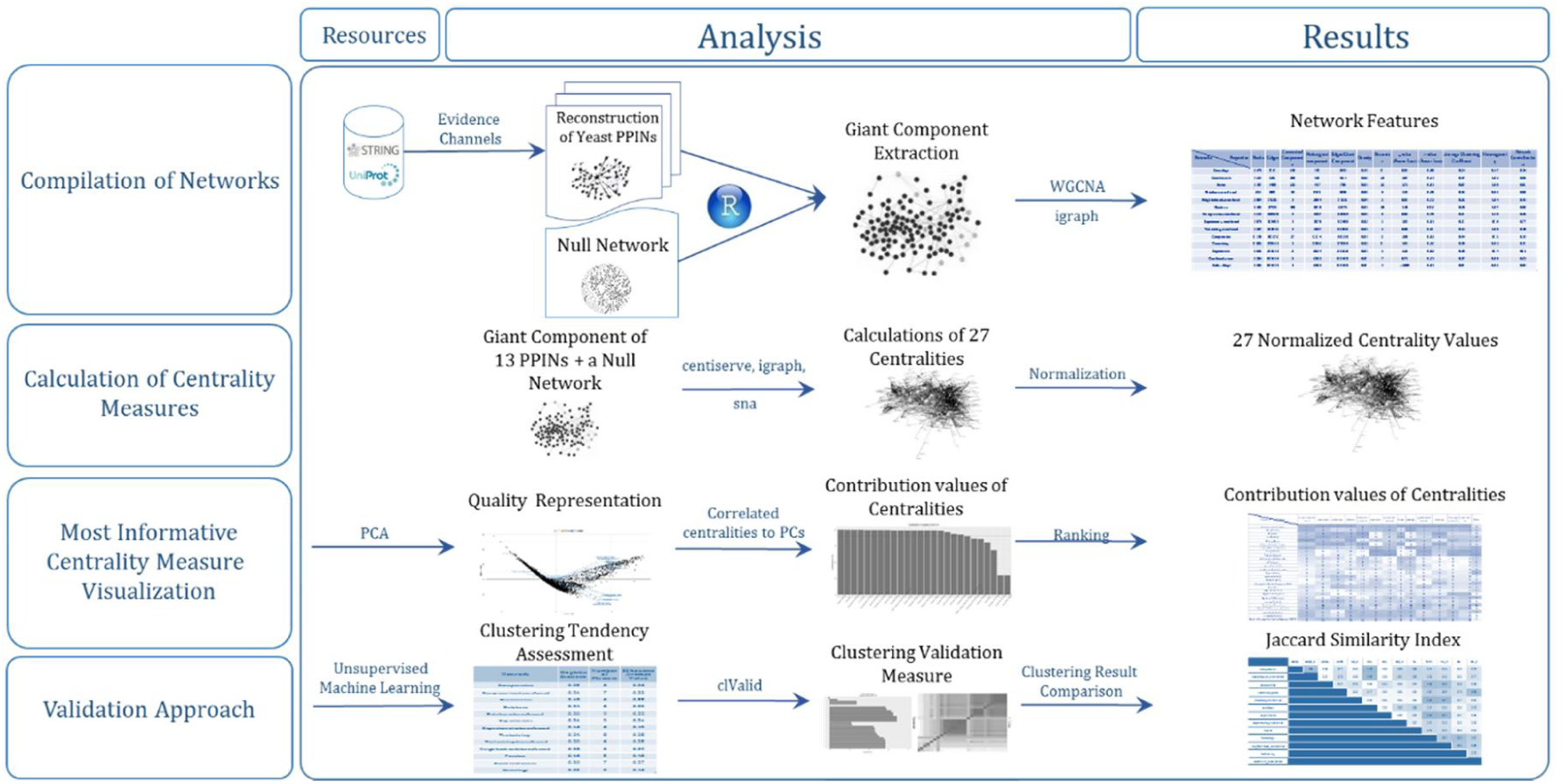
Our workflow for studying the centrality measures. This was followed the reconstruction of the yeast PPIN relying on different kinds of evidence channels as well as the generation of a null network. The workflow contained a comparison of several centrality measures using machine learning methods such as principal components analysis and clustering procedures.

## Reconstruction of the networks

In this study, a UniProtKB reviewed dataset (26) was used to retrieve proteins in *Saccharomyces cerevisiae* (6721 proteins). UniProtKB accessions were converted to STRING using the STRINGdb R package, which resulted in 6603 protein identifiers (3rd Sep 2016). Interactions among proteins were extracted based on the STRING IDs. In the 2017 edition of the STRING database the results of these interactions are structured in a way to provide maximum coverage; this is achieved by including indirect and predicted interactions on the top of the set.(27). In this study, 13 evidence channels (related to the origin and type of evidence) indicating PPIN of yeast were presented: co-expression, co-expression-transferred, co-occurrence, database, database-transferred, experiments, experiments-transferred, fusion, homology, neighborhood-transferred, textmining, textmining-transferred and combined-score. In the following, the name of the reconstructed network is basis of the corresponding channel name which made of. For the purpose of comparison with real network behavior, a null model network was generated. The null network is the Erdos–Rényi model (28) and was generated using the igraph R package (29). The generated null network was created with a size similar to the yeast reconstructed PPIN in order to have a more fair comparison.

## Fundamental network concepts analysis

To understand the network structure, we reviewed various network features using several R packages (30-32). The network density, clustering coefficient, network heterogeneity, and network centralization properties of the network were calculated. The number of connected components and graph diameter for each network were also computed. Then, the power-law distribution was assessed by computing α values and r correlation coefficients. As most of centrality measures require a strongly connected component graph, the giant component of each PPINs and the null network were extracted. Moreover, for a general overview of the structure of the extracted giant components, some network features such as network density, clustering coefficient, network heterogeneity, and network centralization were calculated.

## Centrality analysis

For this research study, we were only considered undirected, loop-free connected graphs according to the PPIN topology. For centrality analysis, the following 27 centrality measures were selected: Average Distance (33), Barycenter (34), Closeness (Freeman) (5), Closeness (Latora) (35), Residual closeness (36), ClusterRank (37), Decay (38), Diffusion degree (39), Density of Maximum Neighborhood Component (DMNC) (40), Geodesic K-Path (41, 42), Katz (43, 44), Laplacian (45), Leverage (46), Lin (47), Lobby (48), Markov (49), Maximum Neighborhood Component (MNC) (40), Radiality (50), Eigenvector (51), Subgraph scores (52), Shortest-Paths betweenness (5), Eccentricity (53), Degree, Kleinberg's authority scores (54), Kleinberg's hub scores (54), Harary graph (53) and Information (55). All these measures may be calculated for undirected networks in a reasonable time. These measures were calculated using the centiserve (7), igraph (29) and sna (56) R packages. For a better visualization, We assorted the centrality measures into five distinct classes including Distance-, Degree-, Eigen-, Neighborhood-based and miscellaneous groups depend on their logic and formulas (Table 1).

**Table 1.** Centrality measures. The centrality measures were represented in five groups depending on their logic and formulae.

| Distance_based | Degree-based | Eigen-based | Neighborhood-based | Miscellaneous |
| --- | --- | --- | --- | --- |
| Average Distance | Authority_score | Eigenvector centralities | ClusterRank | Geodesic K-Path Centrality |
| Barycenter | Degree Centrality | Katz Centrality (Katz Status Index) | Density of Maximum Neighborhood Component (DMNC) | Harary Graph Centrality |
| Closeness Centrality (Freeman) | Diffusion Degree | Laplacian Centrality | Maximum Neighborhood Component (MNC) | Information Centrality |
| Closeness centrality (Latora) | Kleinberg's hub centrality scores |  | Subgraph centrality scores | Markov Centrality |
| Decay Centrality | Leverage Centrality |  |  | Shortest-Paths Betweenness Centrality |
| Eccentricity of the vertices | Lobby Index (Centrality) |  |  |  |
| Lin Centrality |  |  |  |  |
| Radiality Centrality |  |  |  |  |
| Residual Closeness Centrality |  |  |  |  |

## Unsupervised machine learning analysis

Standard normalization (scaling and centering of matrix-like objects) has been undertaken on computed centrality values according to methodology explained in (57). We used PCA, a linear dimensionality reduction algorithm, (58) as a key step to understand which centrality measures better determine central nodes within a network. PCA was done on normalized computed centrality measures. To validate the PCA results in PPINs, we also examined whether the centrality measures in all networks can be clustered according to clustering tendency procedure. To do this, the Hopkins’ statistic values and visualizing VAT (Visual Assessment of cluster Tendency) plots was calculated by factoextra R package (59). We applied the clustering validation measures to access the most appropriate clustering method among hierarchical, kmeans, and PAM (Partitioning Around Medoids) methods using clValid package (60). This provides silhouette scores according to clustering measures which would be helpful for choosing the suitable method. After selection of the clustering technique, factoextra package was used to attain optimal number of clusters (59). In order to measure the dissimilarity among clusters, we used Ward’s minimum variance method. To compare the clustering results in aforementioned PPINs, the Jaccard similarity index was used relying on the similarity metrics of the clustering results within BiRewire package (61).

## Results and Discussion

### Evaluation of network properties

By importing the same set of protein names, the 13 PPINs were extracted from the STRING database using different evidence channels. (Note: the PPI scores derived from the neighborhood channel of yeast were all zero). All these channels distinctly identify an interaction for each protein pair quantitatively. The dependency between evidence channels was also shown in Fig. 2 by a pairwise scatterplot and Pearson’s r correlation coefficient. Most of the networks were not significantly correlated and correlation coefficients were around zero for all networks.

**Fig. 2.**
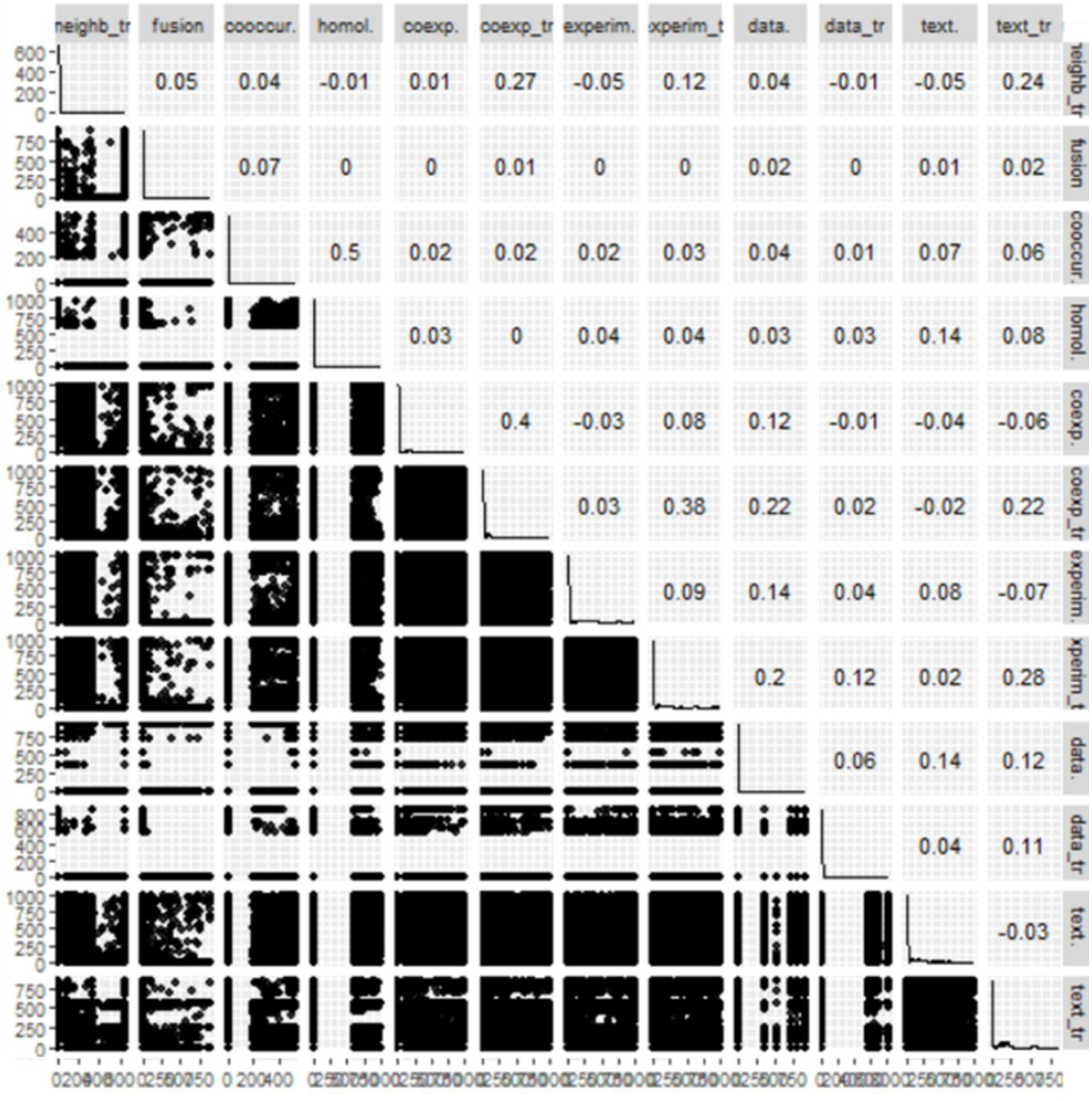
Pairwise scatterplot between the evidence channel scores. The Pearson’s r correlation coefficients between the evidence channels were shown in the upper triangle of the plot. The distributions of scores in each evidence were presented at the diameters of the figure.

In the following, the 14 networks were utilized to undertake an examination of centrality measures. Note that the giant component of each network was accounted for computing several network properties (Table 2). The homology, fusion, co-occurrence and database networks contained high numbers of unconnected components. Except the homology network which had the smallest giant component, the densities of all networks were between 0.01-0.05, as was expected as real network are typically sparse. The network diameter of the fusion, co-occurrence, database and co-expression were one order of magnitude greater than others. All of the PPINs except homology network were correlated to power-law distribution with high r correlation coefficients and diverse alpha power (see Supplementary file 1). The high value of the average clustering coefficients of the database and homology indicated the modular structure of these networks. Compared with the null network, most of the PPINs had a high value of heterogeneity and network centralization. The Degree distribution and clustering coefficients for the networks were also plotted in Fig. 3 and Fig. 4 respectively. Except the homology network, all the Degree distributions were left-skewed similar to scale-free networks. The dependency of PPINs was further assessed and confirmed statistically by Wilcoxon rank sum test (Table 3).

**Fig. 3.**
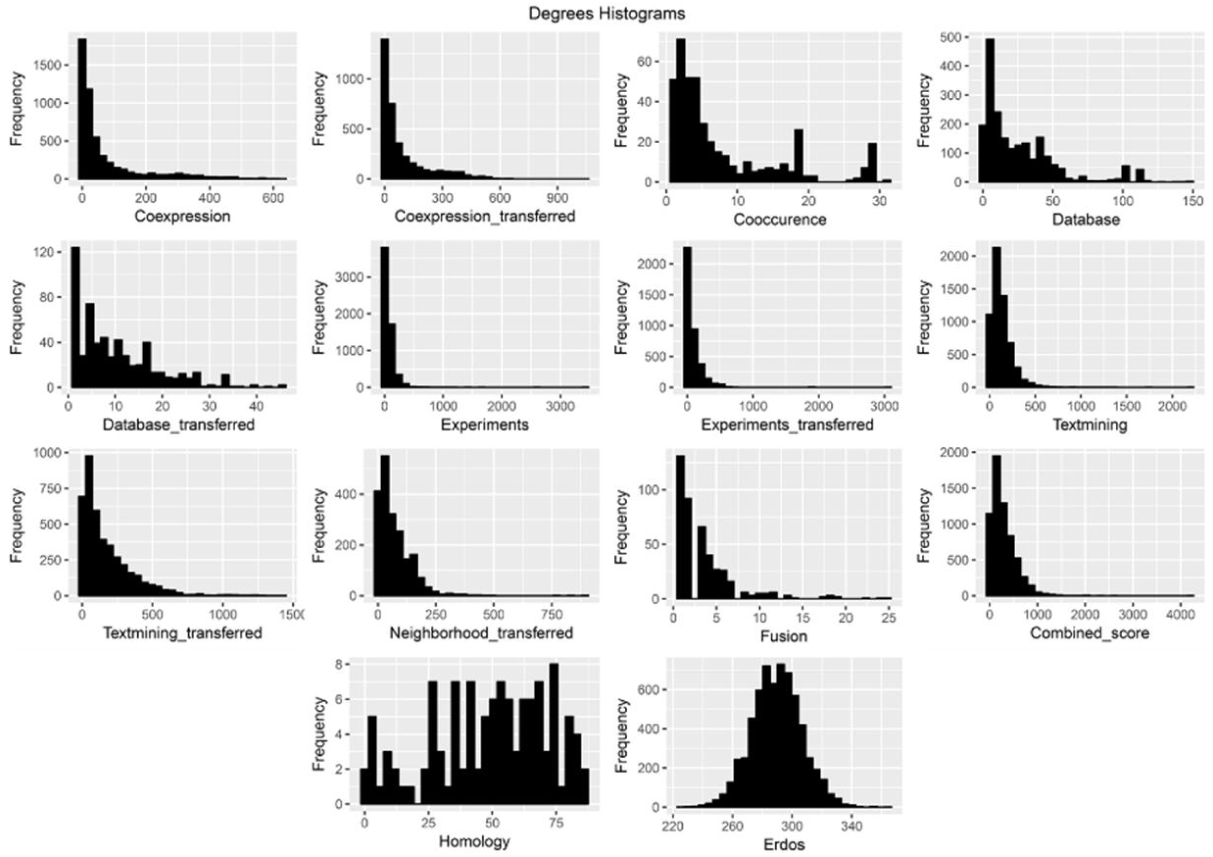
Graphical representation of the Degree distributions in each reconstructed PPIN and the generated null network. The color spectra from green to red indicates low to high values.

**Fig. 4.**
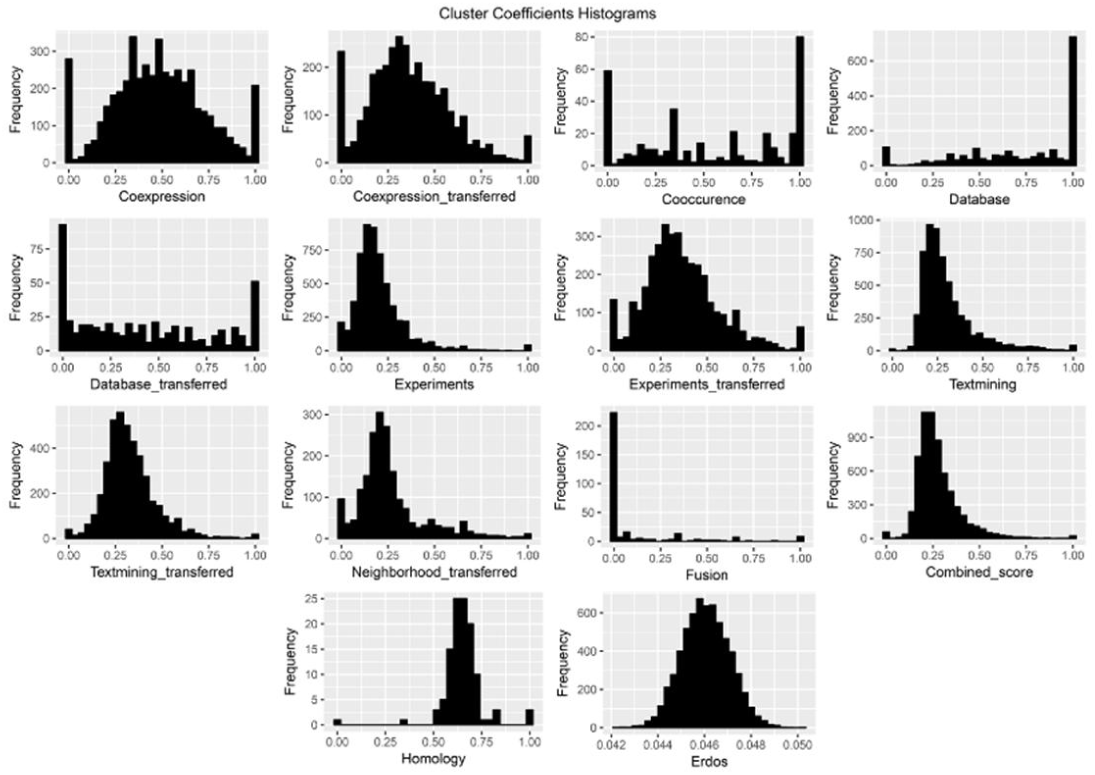
Graphical representation of the clustering coefficient distributions in each reconstructed PPIN and the generated null network.

**Table 2.**
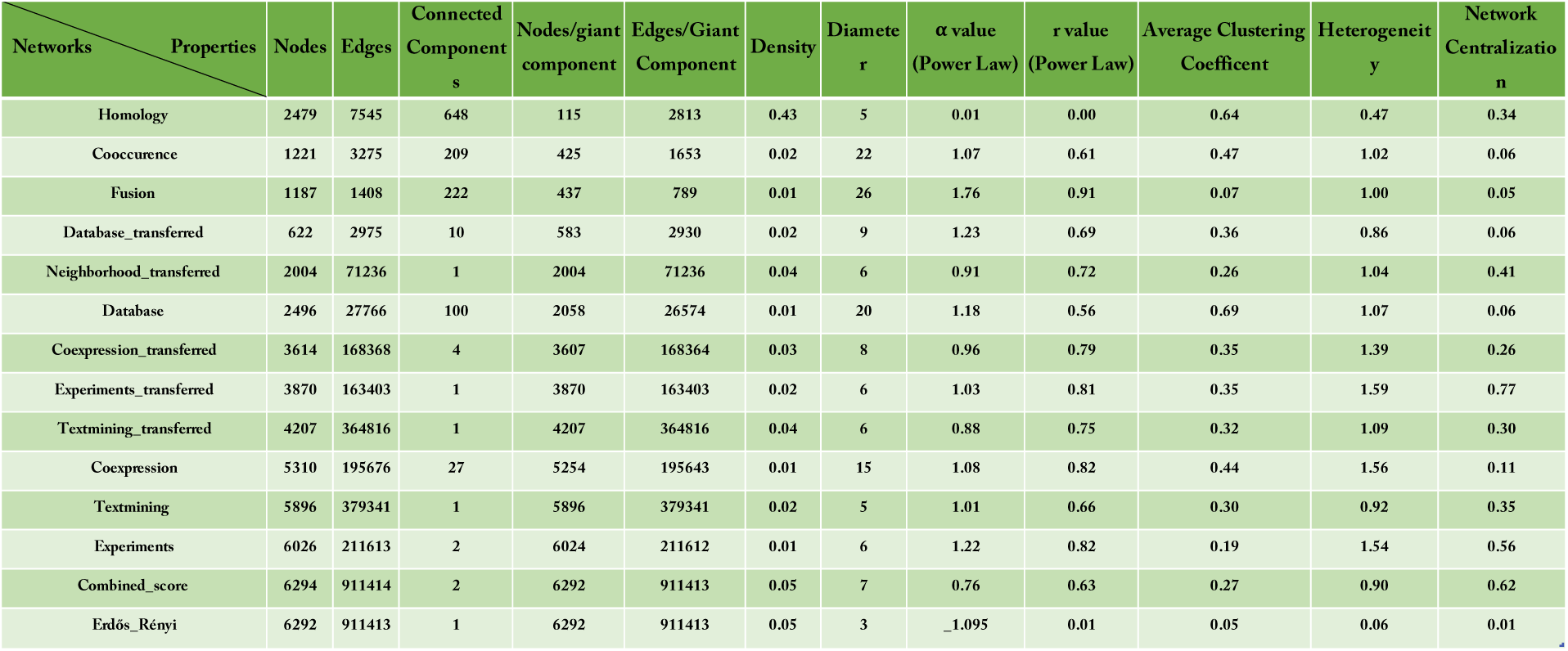
Network global properties of all PPINs and the null network.

**Table 3.** The *p*-value of Wilcoxon rank sum test. The dependency between the distributions of evidence channels evaluated by Wilcoxon test.

|  | coexpressio<br>n | coexpressio<br>n_transfere<br>d | cooccurrence | database | database_tra<br>nsferred | experiments | experiments<br>_transferred | textmining | textmining_t<br>ransferred | neighborhoo<br>d_transfere<br>d | fusion | combined_s<br>core | homology | Erdos |
| --- | --- | --- | --- | --- | --- | --- | --- | --- | --- | --- | --- | --- | --- | --- |
| coexpression |  | 1.02036E-20 | 3.25641E-69 | 2.51433E-23 | 2.78257E-61 | 5.01208E-53 | 5.91091E-38 | < 0.1E+300 | < 0.1E+300 | 6.01743E-46 | 3.6847E-130 | < 0.1E+300 | 3.48803E-05 | < 0.1E+300 |
| coexpression_transferred |  |  | 2.9171E-99 | 7.95167E-73 | 1.18847E-96 | 0.006226156 | 0.016945559 | 2.9123E-186 | 3.2272E-170 | 4.41482E-07 | 4.2326E-159 | < 0.1E+300 | 0.194466705 | < 0.1E+300 |
| cooccurrence |  |  |  | 1.68632E-59 | 1.12261E-06 | 6.9662E-147 | 4.0271E-128 | 1.3673E-228 | 4.0225E-201 | 3.6946E-143 | 7.28332E-20 | 1.522E-229 | 1.05975E-45 | 9.589E-262 |
| database |  |  |  |  | 6.10869E-41 | 8.0539E-163 | 2.931E-118 | < 0.1E+300 | < 0.1E+300 | 1.8392E-144 | 2.3302E-127 | < 0.1E+300 | 1.08534E-22 | < 0.1E+300 |
| database_transferred |  |  |  |  |  | 5.0559E-167 | 1.48E-135 | 1.5733E-290 | 4.4815E-248 | 1.3368E-159 | 1.00973E-45 | 6.4647E-299 | 1.27144E-45 | < 0.1E+300 |
| experiments |  |  |  |  |  |  | 0.914075344 | < 0.1E+300 | 9.1076E-269 | 6.51711E-06 | 9.5956E-200 | < 0.1E+300 | 0.374051268 | < 0.1E+300 |
| experiments_transferred |  |  |  |  |  |  |  | 7.3486E-217 | 9.6268E-187 | 0.000336844 | 1.2341E-184 | < 0.1E+300 | 0.365227158 | < 0.1E+300 |
| textmining |  |  |  |  |  |  |  |  | 1.0304E-07 | 1.3234E-134 | 1.1839E-255 | < 0.1E+300 | 5.39283E-21 | < 0.1E+300 |
| textmining_transferred |  |  |  |  |  |  |  |  |  | 4.7774E-120 | 1.1745E-233 | 2.1904E-166 | 5.63105E-16 | < 0.1E+300 |
| neighborhood_transferred |  |  |  |  |  |  |  |  |  |  | 1.2527E-189 | < 0.1E+300 | 0.277147369 | < 0.1E+300 |
| fusion |  |  |  |  |  |  |  |  |  |  |  | 8.6255E-249 | 4.21561E-53 | 1.337E-268 |
| combined_score |  |  |  |  |  |  |  |  |  |  |  |  | 4.43497E-42 | 6.82527E-90 |
| homology |  |  |  |  |  |  |  |  |  |  |  |  |  | 1.127E-75 |
| erdos |  |  |  |  |  |  |  |  |  |  |  |  |  |  |

### Centrality analysis

In the next step, the 27 centrality measures of nodes were computed in all 14 networks. The distribution and pairwise scatter plots of the computed measures were represented in Fig. 5 to point out pairwise relationship between them. (For the other PPINs see Supplementary file 2). The r correlation coefficients were also shown in this figure in which some of the centrality measures displayed a clear correlation and the others revealed a vast diversity among all five centrality classes. This diversity especially enriched in Distance-, Neighborhood-based and miscellaneous classes for combined-score PPIN compared with Erdos-Renyi network. Analogously, this special profile of centrality measures was repeated in all PPINs to some extent. Another remarkable distinction was the multimodality of distributions in the random network but not in real networks which was repeated for most of the Distance-based centrality measures. Furthermore, according to r correlation coefficients, the pairwise association of centrality measures were roughly higher in the null network than PPINs.

**Fig. 5.**
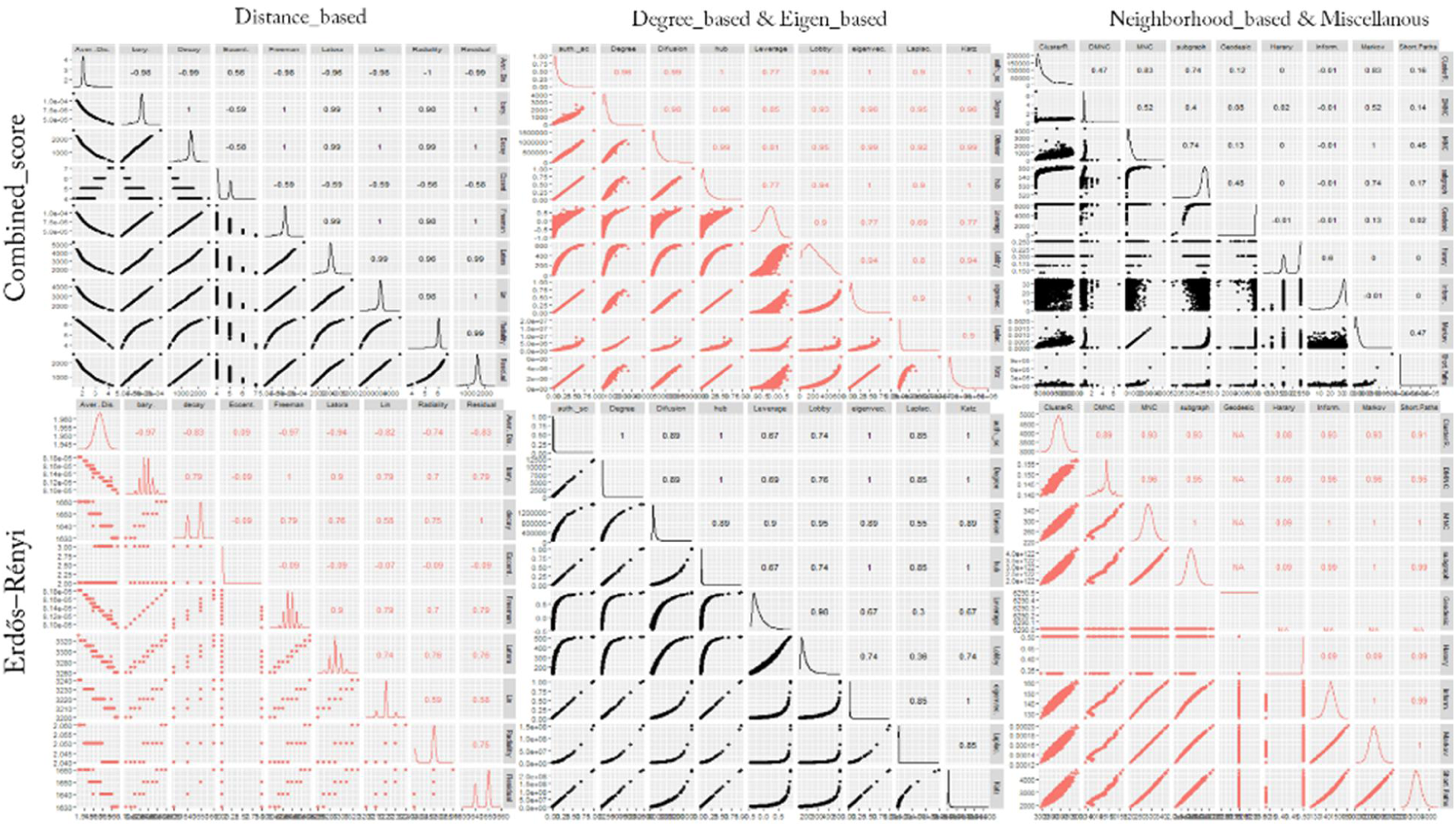
Pairwise scatterplot between the centrality measures. This figure contains combined-score PPIN and the null network. In this figure, the r Pearson correlation coefficients between centralities beside the centralities distribution were also presented in both networks. For better representation, the scatterplot was divided into three parts corresponding to Table 1 groups. For the scatterplot visualizations of all PPINs see Supplementary file 2.

### Dimensionality reduction and clustering analysis

In the next step, PCA-based dimensionality reduction was used to reveal which centrality measures contain the most relevant information in order to effectively identify important or influential nodes in networks. As illustrated in Fig. 6, the profile of the distance to the center of the plot and their directions were mostly consonant except for the homology which was similar to the random network. The rank of contribution values of each centrality measure were shown in Table 4, depend on their corresponding principal components. The percentage of contribution of variables (i.e. centrality measures) in a given PC were computed as (variable.Cos2*100) / (total Cos2 of the component)). A similar profile of the contribution of centrality measures was observed among all biological networks even in homology network opposed to the random null network (See Supplementary file 3). On average, Latora closeness centrality was the major contributor of the principal components in PPINs. In contrast, other well-known centralities i.e. Betweenness and Eccentricity revealed a low contribution value in all PPINs. Analogous to the null network, their values were lower than random threshold depicted in Fig. 8 and Supplementary file 3. On the contrary, the Degree displayed moderate levels of contribution in all real networks whilst it was the fourth rank of random network contributors. Although the profile of contributions were similar, each PPIN exhibited a special fingerprint of the centrality ranking. Finally, by performing unsupervised categorization, we aimed to cluster centrality values computed in the networks. First, we performed a clustering tendency procedure. We found that the centrality values are clusterable in each network as all values in the Hopkins statistics were more than the cutoff (0.05). The results are shown in the first column of Table 5 and Supplementary file 4. Then, by calculating silhouette scores, three methods (i.e. hierarchical, k-means, and PAM) were evaluated in clustering the data sets (Supplementary files 5 & 6). The output of applying these algorithms and the corresponding number of clusters were also shown in Table 5 and Supplementary files 7. Using the hierarchical algorithm based on Ward's method (62), the centrality measures were clustered in each PPINs (Fig. 7). Number of clusters, distance between centrality measures and centrality composition in all 13 PPINs indicated that each centrality ranks nodes within a given network distinctly. For a better comparison, we provided Table 6 containing pairwise Jaccard similarity indices for each network pair. The lowest values were related to the homology, neighborhood-transferred and co-occurrence PPINs while among these genome context prediction methods, fusion PPIN was more associated to the other networks. The high similarity between co-expression and co-expression-transferred was expected however the similar clusters of the database derived PPIN with both aforementioned PPINs and also combined-score with textmining-transferred are noteworthy.

**Fig. 6.**
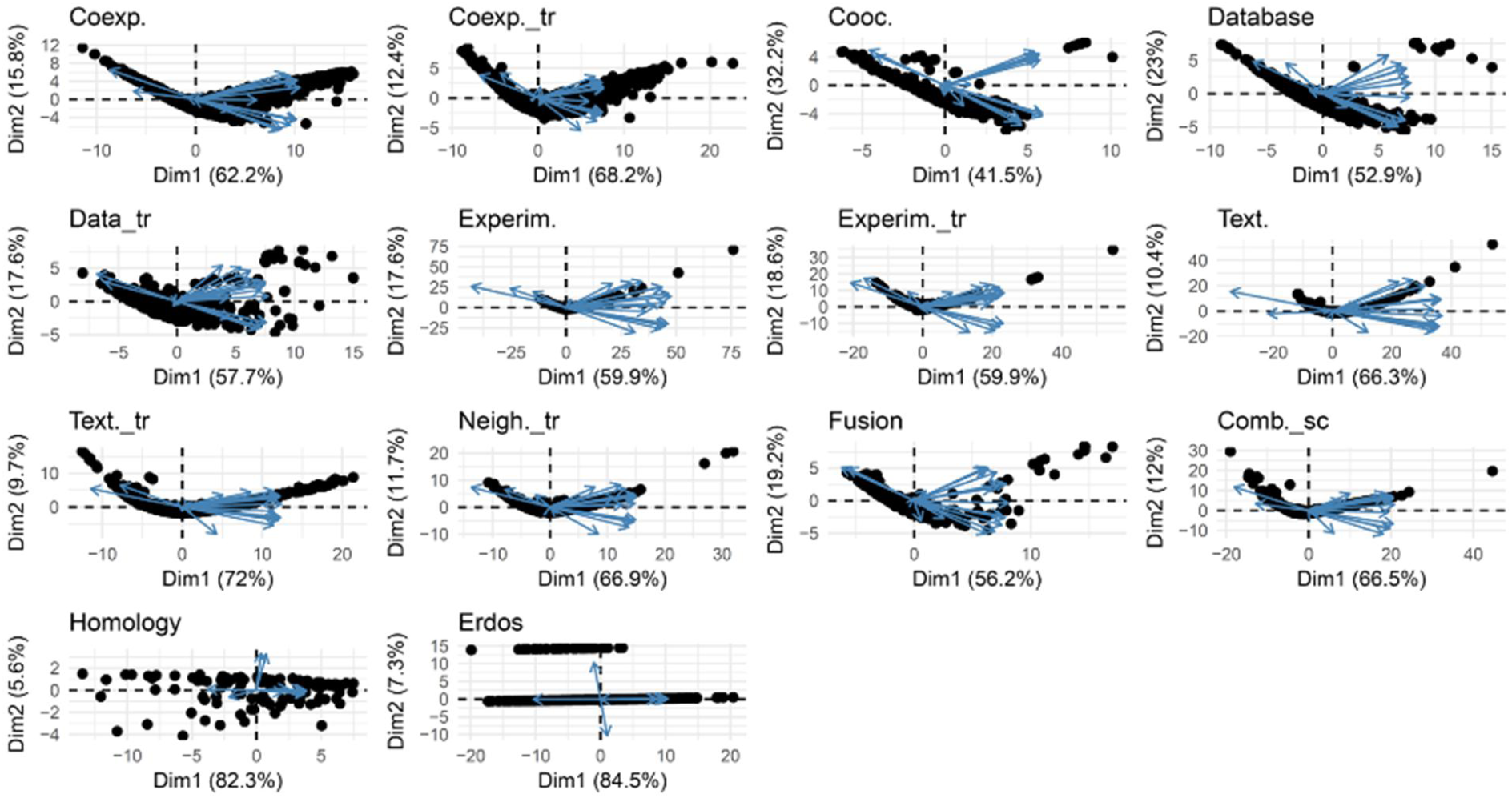
Biplot representation of the centrality measures in each network. The PCA plots were a projection of the multivariate data into the 2D space spanned by the first two principal components. In each plot, nodes were shown as points and centrality measures as vectors.

**Fig. 7.**
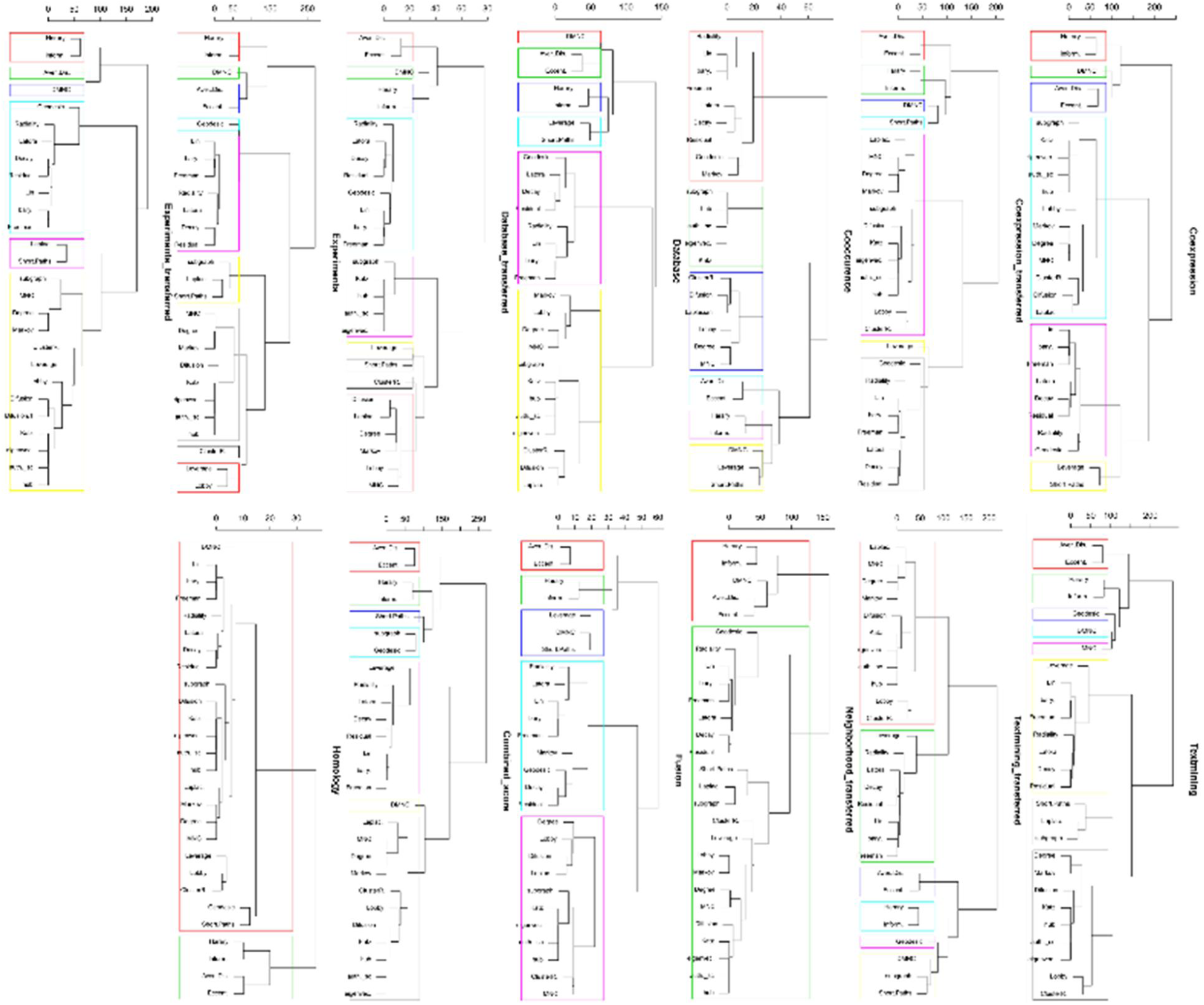
Clustering dendrograms. In each dendrogram, the colored boxes show ensued clusters of centrality measures in each PPIN based on a predefined distance threshold.

**Table 4.**
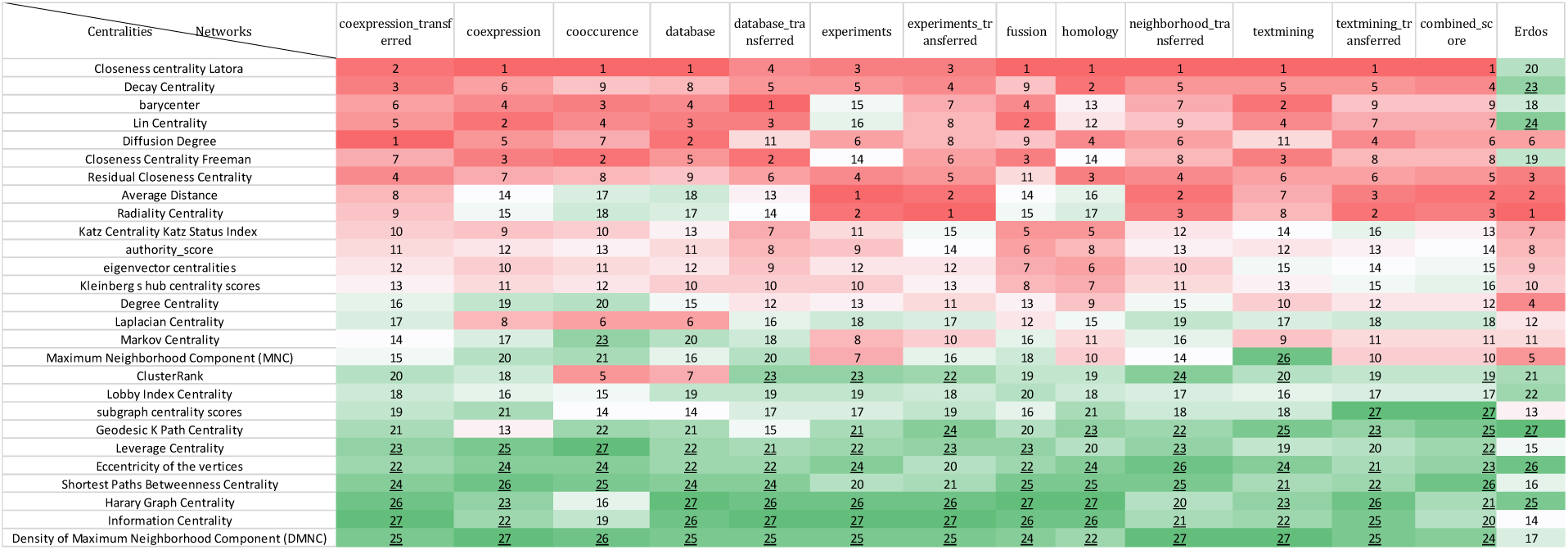
Ranking of the contribution values based on PCA for each network. The red to green highlighted cells represent the top to bottom ranked centrality measures in each network. The underlined ranking values are contribution values of the centrality measures which are below the random threshold.

**Table 5.** Clustering information values for PPINs. The Hopkin’s statistics threshold for clusterability was 0.05.

| Network | Hopkins Statistic | Number of Clusters | Silhouette Average Value |
| --- | --- | --- | --- |
| Coexpression | 0.25 | 6 | 0.36 |
| Coexpression_transferred | 0.21 | 7 | 0.33 |
| Cooccurence | 0.18 | 6 | 0.55 |
| Database | 0.24 | 6 | 0.33 |
| Database_transferred | 0.20 | 9 | 0.32 |
| Experiments | 0.21 | 9 | 0.31 |
| Experiments_transferred | 0.16 | 6 | 0.43 |
| Textmining | 0.24 | 8 | 0.28 |
| Textmining_transferred | 0.20 | 6 | 0.35 |
| Neighborhood_transferred | 0.26 | 2 | 0.39 |
| Fussion | 0.16 | 5 | 0.48 |
| Combined_score | 0.30 | 7 | 0.27 |
| Homology | 0.23 | 2 | 0.46 |

**Table 6.** Jaccard index coefficient values for PPINs. The values represent how similar the networks are, in terms of their clustering results. A value of 1 indicates an exact match while values equal to 0 show dissimilarity.

|  | coexp. | coexp_tr | coocc. | comb. | dat_tr | dat. | exp. | exp_tr | fus. | hom. | nei_tr | tex. | tex_tr |
| --- | --- | --- | --- | --- | --- | --- | --- | --- | --- | --- | --- | --- | --- |
| coexpression |  | 0.99 | 0.58 | 0.77 | 0.62 | 1.00 | 0.58 | 0.80 | 0.83 | 0.41 | 0.43 | 0.62 | 0.76 |
| coexpression_transferred |  |  | 0.57 | 0.78 | 0.63 | 0.99 | 0.58 | 0.81 | 0.82 | 0.40 | 0.43 | 0.62 | 0.77 |
| cooccurrence |  |  |  | 0.47 | 0.75 | 0.58 | 0.44 | 0.50 | 0.73 | 0.29 | 0.30 | 0.43 | 0.48 |
| combined_score |  |  |  |  | 0.52 | 0.77 | 0.62 | 0.63 | 0.64 | 0.37 | 0.39 | 0.78 | 0.96 |
| database_transferred |  |  |  |  |  | 0.62 | 0.55 | 0.55 | 0.55 | 0.25 | 0.27 | 0.47 | 0.51 |
| database |  |  |  |  |  |  | 0.58 | 0.80 | 0.83 | 0.41 | 0.43 | 0.62 | 0.76 |
| experiments |  |  |  |  |  |  |  | 0.59 | 0.49 | 0.25 | 0.27 | 0.67 | 0.63 |
| experiments_transferred |  |  |  |  |  |  |  |  | 0.67 | 0.41 | 0.43 | 0.62 | 0.62 |
| fussion |  |  |  |  |  |  |  |  |  | 0.40 | 0.42 | 0.52 | 0.64 |
| homology |  |  |  |  |  |  |  |  |  |  | 0.91 | 0.30 | 0.37 |
| neighborhood_transferred |  |  |  |  |  |  |  |  |  |  |  | 0.32 | 0.39 |
| textmining |  |  |  |  |  |  |  |  |  |  |  |  | 0.78 |
| textmining_transferred |  |  |  |  |  |  |  |  |  |  |  |  |  |

Interestingly, silhouette scores of centrality measures were closely related to corresponding contribution value of the measures (Fig. 8). Where there was a high silhouette value, a high contribution value was observed, however, a high contribution value did not always mean a high silhouette value. The relationship between the silhouette scores and contribution values of each centrality measure was also examined by regression analysis. Latora closeness, Radiality, Residual, Decay, Lin, Leverage, Freeman closeness and Barycenter centrality measures were present together in the same cluster where the corresponding silhouette scores were all at a high level except the Leverage’s score (Fig. 8A). The average silhouette score was around 0.66 in this cluster. On the other hand, the Leverage’s contribution value was below the threshold line and placed in the group with the least amount of contribution (Fig. 8B). The centrality measures namely Lobby index, ClusterRank, Laplacian, MNC, Degree, Markov, Diffusion degree, Kleinberg's hub, Eigen vector, Authority score, Katz group together where the mean of their silhouette scores (i.e. 0.61) was higher than the overall average and in the same way, their corresponding contribution values were high, too. On the other han, we observed that Shortest path Betweenness (which was in a separated cluster) and Geodesic k path, Subgraph and DMNC (which are all in one cluster) showed the low silhouette value mean (i.e. 0.03) much lower than the average. In all other PPINs, the same relationship between silhouette scores and contribution values was observed as shown in Supplementary files 3 & 6.

**Fig. 8.**
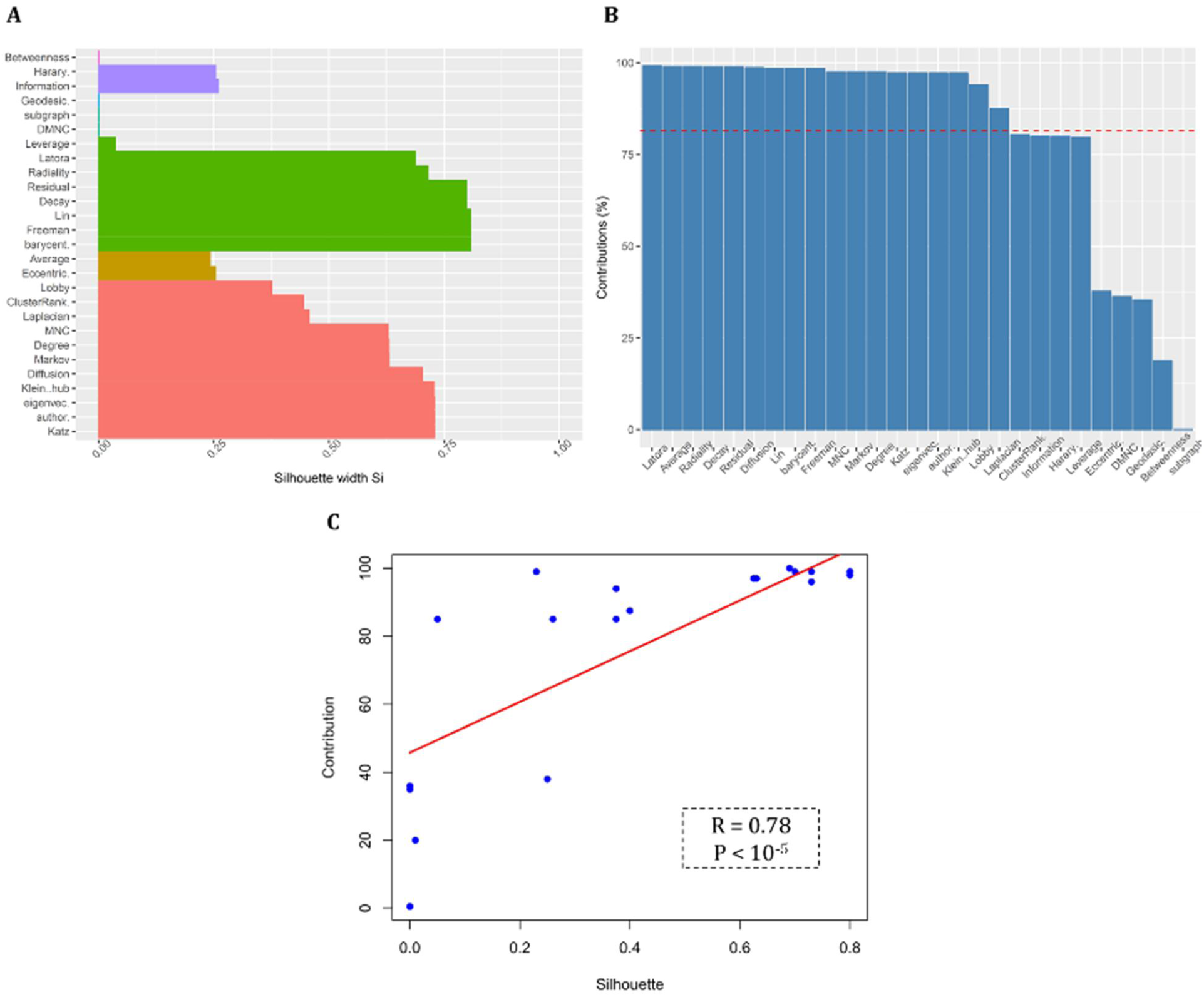
(A) Clustering silhouette plot of the combined-score PPIN. The colors represented the six clusters of the centrality measures in this PPIN. The average silhouette width was 0.49. (B) Contribution values of centrality measures according to their corresponding principal components in this PPIN. The number of principal components stand on the network architecture was equal to 3. The dashed line indicates the random threshold of contribution. (C) Line plot between silhouette and contribution values. The R value shown is the result of a regression coefficient analysis and the p value has been computed from Pearson’s correlation test.

Our results demonstrated that a unique profile of centrality measures including Latora closeness, Barycenter, Diffusion degree, Freeman closeness, Residual, Average distance, Radiality centralities, was the most significant indicator in ranking PPIN nodes. We inferred that the rationale and logic of network reconstruction dictates which centrality measures should be chosen. Also, we demonstrated the relationship between contribution value derived from PCA and silhouette width as a cluster validity index. In this research, we first reasserted that the architecture and global properties of a network impact on the central component detection and that the center of a network would be different, depending on the network’s inherent topology. In other words, we addressed this issue whether a given centrality measure has enough information via-a-vis and it demonstrates a same behavior in some other networks.

## Conclusion

Network-based methods have been introduced as an emergent approach for simplification, reconstruction, analysis, and comprehension of complex behavior in biological systems. Network-based ranking methods (i.e. centrality analysis) have been found widespread use for predicting essential proteins, proposing drug targets candidates in treatment of cancer, biomarker discovery, human disease genes identification and creation a cell with the minimal genome (63). However, there is no consensus pipeline for centrality analysis regarding aforementioned applications among network analysts.

In this study, we worked on yeast PPINs which were built using 13 evidence channels in the STRING database. Subsequently, 27 centrality measures were used for the prioritization of the nodes in all PPINs. We illustrated that data reduction and low-dimensional projection help to extract relevant features (i.e. centrality measures) and corresponding relationships. Thus, to quantify connectivity in biological networks, we recommend that before calculating centrality measures, PCA (as an example of data projection methods) conduce how to use these measures. In the other word, the analysis of principal components clarifies which measures have the highest contribution values, i.e., which measures comprise much more information about centrality.

## Basic definitions

- **Giant component of a graph** defines the largest connected component of a graph in which there is a path between each pair of nodes (64).
- **Network density** is a representation of the number of interactions to the number of possible interactions among a given network (65).
- **Network centralization** refers to a topological spectrum from star to grid topologies (where each node has a same number of links) of a graph varies from 1 to 0 (65).
- The **network heterogeneity** measure describes as the coefficient of variation of connectivity distribution. A high heterogeneous network implies that the network is exhibited approximate scale-free topology (65).
- The **clustering coefficient** of a node is the number of triangles (3-loops) that pass through it, relative to the maximum number of 3-loops that could pass through the node. The network clustering coefficient defines as the mean of the clustering coefficients for all nodes in the network (66).
- **Influential nodes** which is generally used in social networks analysis point as nodes with good spreading properties in networks (67). Different centrality measures are used to find influential nodes.
- **Centrality-lethality rule** explains nodes with high centrality values in which maintain the integrity of the network structure, are more related to the biological system survival (68).
- The **silhouette criterion** defines how similar a centrality is to its own cluster compared to other clusters. It ranges from -1 to 1, where a high value infers that the centrality is well matched to its own cluster and poorly matched to neighboring clusters. If most centralities have a high value, then the clustering configuration is proper. If they have low or negative values, then the clustering configuration may have too many or too few clusters (69).
- In order to see definitions of all used centrality measures, see http://www.centiserver.org.

## Supplementary data

Supplementary file 1: Fitted power law distribution. The Degree distribution of each network has been compared to the power law distribution in order to visualize the scale free property in the structure of each network.

Supplementary file 2: Scatterplots between groups of centralities. Each panel indicates scatterplots between centralities groups of two networks.

Supplementary file 3: Contribution values of centralities in each network. These values were computed based on the principal components. The red line shows the threshold used for identifying effective centralities.

Supplementary file 4: Visual assessment of cluster tendency plots. Each rectangular represents the clusters of the calculated results of the centrality measures.

Supplementary file 5: Clustering properties results. These properties include connectivity, Dunn and Silhouette scores. These scores suggest the sufficient clustering method by a specific number of clusters.

Supplementary file 6: Clusters silhouette plots. Each color represents a cluster and each bar with specific color indicates a centrality.

Supplementary file 7: Optimal number of clusters. The suitable number of clusters for hierarchical clustering method was computed using the average silhouette values.

## References

1. Jeong H, Mason SP, Barabasi A-L, Oltvai ZN. Lethality and centrality in protein networks. arXiv preprint cond-mat/0105306. 2001.

2. Csermely P, Korcsmáros T, Kiss HJ, London G, Nussinov R. Structure and dynamics of molecular networks: a novel paradigm of drug discovery: a comprehensive review. Pharmacology & therapeutics. 2013;138(3):333–408.

3. Freeman LC. Going the Wrong Way on a One-Way Street: Centrality in Physics and Biology. Journal of Social Structure. 2008;9(2):1–15.

4. Landherr A, Friedl B, Heidemann J. A Critical Review of Centrality Measures in Social Networks. Bus Inform Syst Eng+. 2010;2(6):371–85.

5. Freeman LC. Centrality in social networks conceptual clarification. Social networks. 1978;1(3):215–39.

6. Buttner K, Scheffler K, Czycholl I, Krieter J. Social network analysis - centrality parameters and individual network positions of agonistic behavior in pigs over three different age levels. Springerplus. 2015;4:185.

7. Jalili M, Salehzadeh-Yazdi A, Asgari Y, Arab SS, Yaghmaie M, Ghavamzadeh A, et al. CentiServer: A Comprehensive Resource, Web-Based Application and R Package for Centrality Analysis. PLoS one. 2015;10(11):e0143111.

8. Hahn MW, Kern AD. Comparative genomics of centrality and essentiality in three eukaryotic protein-interaction networks. Molecular Biology and Evolution. 2005;22(4):803–6.

9. Bergmann S, Ihmels J, Barkai N. Similarities and differences in genome-wide expression data of six organisms. PLoS biology. 2004;2(1):E9–E.

10. Wagner A, Fell DA. The small world inside large metabolic networks. Proceedings Biological sciences / The Royal Society. 2001;268(1478):1803–10.

11. Ma HW, Zeng AP. The connectivity structure, giant strong component and centrality of metabolic networks. Bioinformatics. 2003;19(11):1423–30.

12. Estrada E. Virtual identification of essential proteins within the protein interaction network of yeast. Proteomics. 2006;6(1):35–40.

13. He X, Zhang J. Why do hubs tend to be essential in protein networks? PLoS genetics. 2006;2(6):e88–e.

14. Joy MP, Brock A, Ingber DE, Huang S. High-betweenness proteins in the yeast protein interaction network. Journal of Biomedicine and Biotechnology. 2005;2005(2):96–103.

15. Potapov AP, Voss N, Sasse N, Wingender E. Topology of mammalian transcription networks. Genome informatics International Conference on Genome Informatics. 2005;16(2):270–8.

16. Yang Y, Han L, Yuan Y, Li J, Hei N, Liang H. Gene co-expression network analysis reveals common system-level properties of prognostic genes across cancer types. Nature communications. 2014;5.

17. Tew KL, Li XL, Tan SH. Functional centrality: Detecting lethality of proteins in protein interaction networks. Genome Inform Ser. 2007;19:166–77.

18. Peng X, Wang J, Wang J, Wu FX, Pan Y. Rechecking the Centrality-Lethality Rule in the Scope of Protein Subcellular Localization Interaction Networks. PLoS One. 2015;10(6):e0130743.

19. Khuri S, Wuchty S. Essentiality and centrality in protein interaction networks revisited. BMC bioinformatics. 2015;16(1):1.

20. Koschützki D, Schreiber F, editors. Comparison of Centralities for Biological Networks. German Conference on Bioinformatics; 2004: Citeseer.

21. Dwyer T, Hong S-H, Koschützki D, Schreiber F, Xu K, editors. Visual analysis of network centralities. Proceedings of the 2006 Asia-Pacific Symposium on Information Visualisation-Volume 60; 2006: Australian Computer Society, Inc.

22. Valente TW, Coronges K, Lakon C, Costenbader E. How correlated are network centrality measures? Connections (Toronto, Ont). 2008;28(1):16.

23. Batool K, Niazi MA. Towards a Methodology for Validation of Centrality Measures in Complex Networks (vol 9, e90283, 2014). PLoS One. 2014;9(5).

24. Li C, Li Q, Van Mieghem P, Stanley HE, Wang H. Correlation between centrality metrics and their application to the opinion model. The European Physical Journal B. 2015;88(3):65.

25. Jalili M, Salehzadeh-Yazdi A, Gupta S, Wolkenhauer O, Yaghmaie M, Resendis-Antonio O, et al. Evolution of Centrality Measurements for the Detection of Essential Proteins in Biological Networks. Front Physiol. 2016;7:375.

26. Boutet E, Lieberherr D, Tognolli M, Schneider M, Bansal P, Bridge AJ, et al. UniProtKB/Swiss-Prot, the manually annotated section of the UniProt KnowledgeBase: how to use the entry view. Plant Bioinformatics: Methods and Protocols. 2016:23–54.

27. Szklarczyk D, Morris JH, Cook H, Kuhn M, Wyder S, Simonovic M, et al. The STRING database in 2017: quality-controlled protein–protein association networks, made broadly accessible. Nucleic Acids Research. 2016:gkw937.

28. Erdos P, Rényi A. On the evolution of random graphs. Publ Math Inst Hung Acad Sci. 1960;5(1):17–60.

29. Csardi G, Nepusz T. The igraph software package for complex network research. InterJournal, Complex Systems. 2006;1695(5):1–9.

30. Butts CT. network: a Package for Managing Relational Data in R. Journal of Statistical Software. 2008;24(2):1–36.

31. Langfelder P, Horvath S. WGCNA: an R package for weighted correlation network analysis. BMC bioinformatics. 2008;9(1):1.

32. Horvath S. Weighted network analysis: applications in genomics and systems biology:Springer Science & Business Media; 2011.

33. del Rio G, Koschützki D, Coello G. How to identify essential genes from molecular networks? BMC Systems Biology. 2009;3(1):1.

34. Viswanath M. Ontology-based automatic text summarization. University of Georgia. 2009.

35. Latora V, Marchiori M. Efficient behavior of small-world networks. Physical review letters. 2001;87(19):198701.

36. Dangalchev C. Residual closeness in networks. Physica A: Statistical Mechanics and its Applications. 2006;365(2):556–64.

37. Chen D-B, Gao H, Lü L, Zhou T. Identifying influential nodes in large-scale directed networks: the role of clustering. PLoS one. 2013;8(10):e77455.

38. Hurajová J, Gago S, Madaras T. Decay Centrality. 15th Conference of Košice Mathematicians; 2.–5. apríla; Herl'any2014.

39. Kundu S, Murthy CA, Pal SK. A New Centrality Measure for Influence Maximization in Social Networks. Lect Notes Comput Sc. 2011;6744:242–7.

40. Lin C-Y, Chin C-H, Wu H-H, Chen S-H, Ho C-W, Ko M-T. Hubba: hub objects analyzer—a framework of interactome hubs identification for network biology. Nucleic acids research. 2008;36(Suppl 2):W438–W43.

41. Borgatti SP, Everett MG. A graph-theoretic perspective on centrality. Social networks. 2006;28(4):466–84.

42. De Meo P, Ferrara E, Fiumara G, Ricciardello A. A novel measure of edge centrality in social networks. Knowledge-based systems. 2012;30:136–50.

43. Grassler J, Koschutzki D, Schreiber F. CentiLib: comprehensive analysis and exploration of network centralities. Bioinformatics. 2012;28(8):1178–9.

44. Junker BH, Koschützki D, Schreiber F. Exploration of biological network centralities with CentiBiN. BMC bioinformatics. 2006;7(1):1.

45. Qi X, Fuller E, Wu Q, Wu Y, Zhang C-Q. Laplacian centrality: A new centrality measure for weighted networks. Information Sciences. 2012;194:240–53.

46. Joyce KE, Laurienti PJ, Burdette JH, Hayasaka S. A new measure of centrality for brain networks. PLoS One. 2010;5(8):e12200.

47. Hoffman AN, Stearns TM, Shrader CB. Structure, context, and centrality in interorganizational networks. Journal of Business Research. 1990;20(4):333–47.

48. Korn A, Schubert A, Telcs A. Lobby index in networks. Physica A: Statistical Mechanics and its Applications. 2009;388(11):2221–6.

49. White S, Smyth P, editors. Algorithms for estimating relative importance in networks. Proceedings of the ninth ACM SIGKDD international conference on Knowledge discovery and data mining; 2003: ACM.

50. Zotenko E, Mestre J, O'Leary DP, Przytycka TM. Why do hubs in the yeast protein interaction network tend to be essential: reexamining the connection between the network topology and essentiality. PLoS Comput Biol. 2008;4(8):e1000140.

51. Bonacich P. Power and centrality: A family of measures. American journal of sociology. 1987:1170–82.

52. Estrada E, Rodriguez-Velazquez JA. Subgraph centrality in complex networks. Physical Review E. 2005;71(5):056103.

53. Hage P, Harary F. Eccentricity and centrality in networks. Social networks. 1995;17(1):57–63.

54. Kleinberg JM. Authoritative sources in a hyperlinked environment. Journal of the ACM (JACM). 1999;46(5):604–32.

55. Stephenson K, Zelen M. Rethinking centrality: Methods and examples. Social Networks. 1989;11(1):1–37.

56. Butts CT. sna: Tools for social network analysis. R package version. 2010;2(2).

57. Becker RA, Chambers JM, Wilks AR. The new S language. Pacific Grove, Ca: Wadsworth & Brooks, 1988. 1988.

58. Abdi H, Williams LJ. Principal component analysis. Wiley Interdisciplinary Reviews:Computational Statistics. 2010;2(4):433–59.

59. Kassambara. A. factoextra: Visualization of the outputs of a multivariate analysis. R Package version 1.0. 1. 2015.

60. Brock G, Pihur V, Datta S, Datta S. clValid, an R package for cluster validation. Journal of Statistical Software (Brock et al, March 2008). 2011.

61. Gobbi A, Albanese D, Iorio F. Package 'BiRewire’. 2016.

62. Ward Jr JH. Hierarchical grouping to optimize an objective function. Journal of the American statistical association. 1963;58(301):236–44.

63. Glass JI, Hutchison CA, 3rd, Smith HO, Venter JC. A systems biology tour de force for a near-minimal bacterium. Mol Syst Biol. 2009;5:330.

64. Barneh F, Jafari M, Mirzaie M. Updates on drug-target network; facilitating polypharmacology and data integration by growth of DrugBank database. Briefings in bioinformatics. 2016;17(6):1070–80.

65. Horvath S. Weighted Network Analysis. New York, NY: Springer New York; 2011.

66. Analysis of Biological Networks. Hoboken, NJ, USA: John Wiley & Sons, Inc.; 2008.

67. Malliaros FD, Rossi M-EG, Vazirgiannis M. Locating influential nodes in complex networks. Scientific Reports. 2016;6(1):19307-.

68. Jeong H, Mason SP, Barabasi AL, Oltvai ZN. Lethality and centrality in protein networks. Nature. 2001;411(6833):41–2.

69. Rousseeuw PJ. Silhouettes: A graphical aid to the interpretation and validation of cluster analysis. Journal of Computational and Applied Mathematics. 1987;20:53–65.

